# Dissecting heterogeneous brain development and aging using voxelwise normative models

**DOI:** 10.64898/2026.03.27.714738

**Authors:** Alice V. Chavanne, Yaping Wang, Augustijn A. A. de Boer, Bing Xu, Teije H. van Prooije, Kirsten C. J. Kapteijns, Colette Reniers, Carlos R. Hernandez-Castillo, Juan Fernandez-Ruiz, Bart P. van de Warrenburg, Jörn Diedrichsen, Ryan L. Muetzel, André F. Marquand

## Abstract

Brain disorders are often characterized by biological heterogeneity that is poorly captured by group-average analyses. Normative modeling has emerged as a promising tool to parse out such heterogeneity, yet existing lifespan reference models rely on coarse parcellations, which may obscure individual variability. Using an aggregated reference sample (n=58,597 scans from n=51,107 participants), we provide openly available normative models of brain morphometry at the voxel level across the lifespan, and we illustrate their potential utility with two complementary applications. First, we investigated long-term brain development after preterm birth across two independent cohorts (n=284; n=304) and found individualized, replicable and persistent brain alterations. Second, we extracted high-resolution patient-level morphometric deviations in two samples with rare, genetic neurodegenerative disorders (spinocerebellar ataxia type 1 and 3; n=29, n=15), which showed marked heterogeneity. Together, our findings highlight that voxelwise normative modeling can detect clinically relevant, individualized deviations from a reference model with high spatial precision.

## Introduction

Neurological and psychiatric disorders together represent the largest global burden of disease, as well as heavy societal and economic costs in- and outside of healthcare^1–3^. Such disorders are often characterized by clinical and biological heterogeneity, hindering effective diagnosis and treatment allocation^4^. While brain disorders have been associated with atypical anatomical changes^5,6^, healthy age-related changes in the brain also show considerable inter-individual variability^7^.

The emergence of normative modeling offers a promising, person-centered tool to move beyond case-control analyses and parse such biological heterogeneity by leveraging existing large neuroimaging datasets^8–10^. This approach was designed to chart the trajectory of biological variables across covariates of interest (e.g. age) in aggregated large-scale samples, and enables the calculation of deviations from that population reference norm at the individual level. Importantly, it enables the calculation of normative deviations in unseen samples, provided a calibration subsample is used to account for new site effects.

Many large-scale normative reference models have been made available online^7,11–14^ and normative models have successfully been used to shed light on the neuroanatomical heterogeneity of many disorders, such as schizophrenia, bipolar disorder, attention deficit hyperactivity disorder, autism and Alzheimer’s disease^15–18^. However, these reference models and studies have nearly exclusively examined normative deviations at the level of relatively coarse brain regions. This is important because region-level models obscure inter-individual variability and lack sensitivity to capture more subtle or localized effects on neuroanatomy. To date, only two lifespan voxel-level volumetric normative models have been published, which were bespoke models designed to examine deviations in childhood adversity or copy number variant carriers, respectively^19,20^. These were also based on a brain image preprocessing pipeline that has been shown to yield inferior performance compared to other alternatives^21^.

To address this need, we present voxelwise normative models of brain development and aging across the lifespan estimated with a more performant MRI preprocessing pipeline and an aggregated large-scale sample. These models, which chart age-related trajectories at unprecedented spatial precision, allow for more comprehensive characterization of brain morphometry that has been lacking for granular individual-level inference within the normative modeling framework. We demonstrate the potential utility of our models using two complementary use cases. First, we illustrate the heterogeneity in long-term brain development after preterm birth using two independent longitudinal cohorts, spanning early childhood to adolescence. Second, we produce patient-level, whole-brain maps of extreme normative deviations in two variants of a rare genetically inherited neurodegenerative condition, spinocerebellar ataxia type 1 and 3 (SCA1 and SCA3), from two independent samples, and conduct exploratory predictions of disease severity and disease progression using normative measures.

Preterm birth, defined as < 37 weeks of gestation, occurs in more than 10% of births worldwide and is the leading cause of perinatal mortality^22^. It has also been associated with an elevated risk of long-term neurological and neurodevelopmental impairments^23,24^.While these risks are generally more severe in individuals born very prematurely, neurodevelopmental trajectories have been shown to vary significantly across individuals born with the same term duration^25,26^. Previous studies have identified altered brain anatomy in widespread areas in preterm infants compared to individuals born-at-term, but they have also shown substantial neuroanatomical heterogeneity across individuals born preterm, with non-overlapping alteration patterns^27–29^. A recent study investigated long-term effects of preterm birth on brain anatomy using a normative modeling approach, but only cortical, region-level variables were examined^30^.

Spinocerebellar ataxias (SCA) are rare, autosomal dominant neurodegenerative diseases, with a prevalence of approximately 3 affected individuals per 100,000 individuals^31^. SCA1 and SCA3 are among the most common SCA subtypes and are caused by a CAG repeat expansion in the *ATXN1* and *ATXN3* genes, respectively. Both subtypes are characterized by cerebellar ataxia, causing symptoms such as unsteady gait, impaired coordination of limbs and slow speech, although non-ataxia symptoms are also frequent. SCA1 and SCA3 lead to severe disability and a substantially reduced life expectancy^32^. However, no disease-modifying treatment is available for either SCA subtype to date and, given the rarity of these diseases, sample sizes estimates for potential clinical trials are prohibitive^33^. Thus, biomarkers of treatment effects and disease progression are needed to reduce sample size requirements, with structural neuroimaging being one of the promising avenues to detect candidate markers for SCA. Neuroimaging findings in SCA1 and SCA3 have shown broad overlap, including marked cerebellar and pontine atrophy^31^, and regional volumes in the brainstem and cerebellum have been associated with prospective disease progression in SCA1^34^. However, clinical and neuroanatomical heterogeneity across patients remains, that cannot be tackled by group-level analyses.

These complementary use cases, a common birth outcome and a rare neurodegenerative disorder, underscore the necessity of characterizing interindividual differences across the lifespan at high spatial precision and with whole-brain coverage, moving beyond coarse regional estimates. They demonstrate the utility of voxelwise normative models in identifying nuanced, fine-grained and individualized patterns of atypicality, providing a more complete perspective of the heterogeneity underlying diverse clinical conditions.

## Methods

Please see Fig.1 for a general presentation of the workflow. We made all the code used in this study openly available at https://github.com/RaynorCerebellumCharts/NormativeModels. The estimated normative models are available at (https://surfdrive.surf.nl/s/Mb6mZyFmJeCaPcZ).

**Figure 1:**
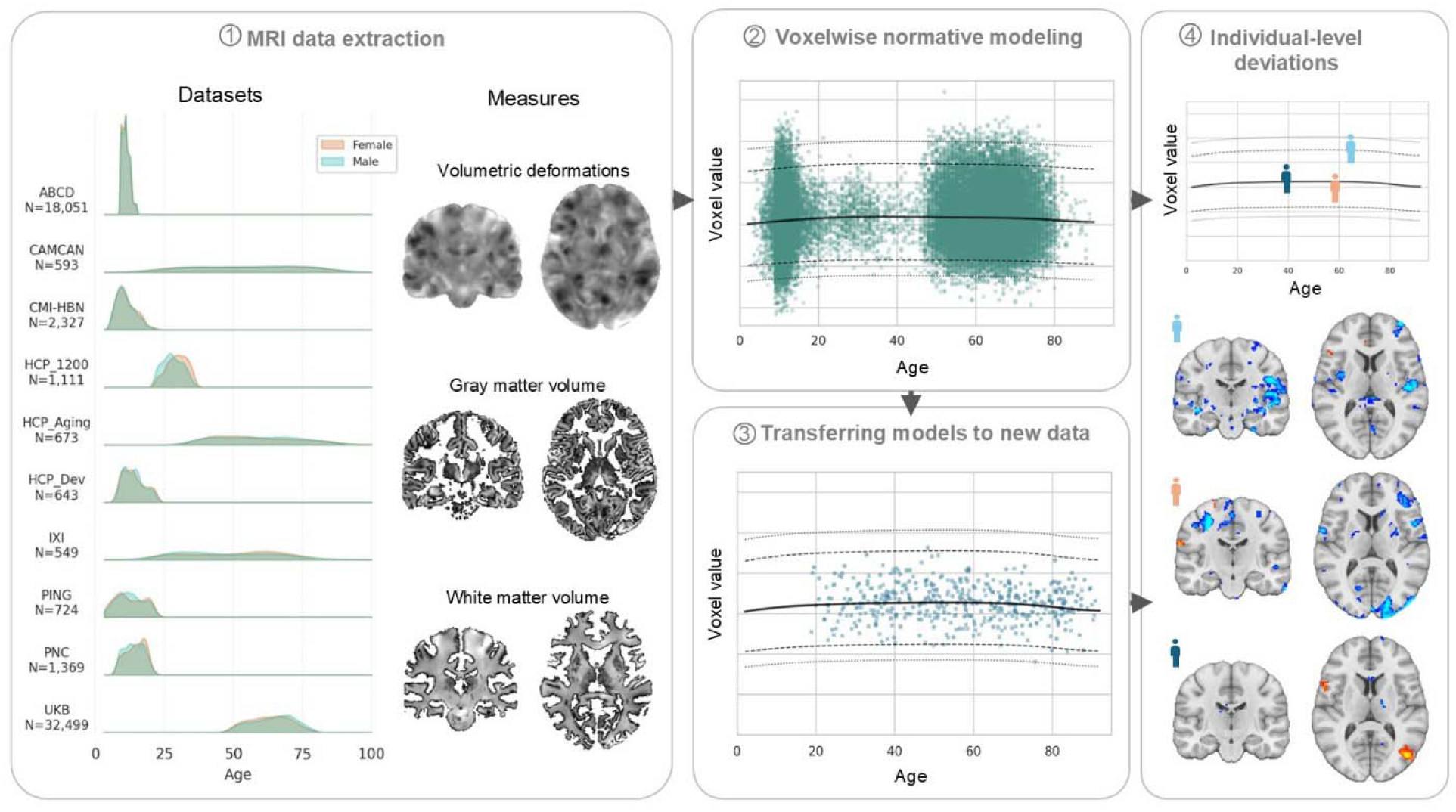
Overview of datasets and analysis pipeline. Using an aggregated sample of 10 datasets, voxelwise normative models were estimated from structural MRI measures: log jacobian determinants (n=58,539 quality-controlled scans), grey matter volume and white matter volume (n=53,593 quality-controlled scans for both). Estimated models were transferred to unseen samples using a calibration subset. Finally, individual-level voxelwise maps of extranormal deviations were extracted for participants of interest.

### 1. Datasets, preprocessing and brain measures extraction

We aggregated a large-scale reference samples from from the UK Biobank^35^, Cam-CAN^36^, PNC^37^, HCP-Aging^38^, HCP-Dev^39^, HCP-1200^40^, ABCD^41,42^(release 4.0), IXI (https://brain-development.org/ixi-dataset/), PING^43^, and CMI-HBN^44^. MRI acquisition parameters are described in the respective studies.

We preprocessed all T1-weighted anatomical 3T MRI scans using the ANTs module (version 2.2, except for ABCD and CMI-HBN that were processed with version 2.5.4), including default N4 bias correction, brain extraction, and Syn registration (rigid, affine and non-linear) to standard space using the MNI152NLin2009cSym template (1x1x1 mm^3^ isotropic resolution). We visually inspected registered T1-weighted scans for artefacts, excessive movement, misregistration and pipeline failures (N=3,290 scans excluded).

Jacobian determinants (JDs) of the non-linear deformation field measure, the degree of volume contraction or expansion required at each voxel to fit subject data to the template. The use of JDs prevents potentially arbitrary distinctions between grey and white matter, avoiding some partial volume effects. Thus, log(JDs) from combined affine and nonlinear transformations (calculated with the geometric method in ANTs) were chosen as a voxelwise measure of subject-level brain anatomy (for further details see ^19,20^).

We also extracted modulated grey and white matter volumes (GMV and WMV), as additional voxelwise measures of brain anatomy. To this end, we processed all MRI scans in FSL (version 6.0.6) with the default FAST step from the *fsl_anat* pipeline, including image reorientation to standard, automatic cropping, bias-field correction and tissue segmentation. We applied the registration transforms previously calculated with ANTs to the segmented grey and white matter densities from FAST. We visually inspected registered tissue densities for brain extraction and segmentation failures, and we modulated valid images with the JDs to obtain GMV and WMV.

We excluded scan sites with < 60 valid observations to allow for accurate estimation of site/scanner effects. Final sample sizes with available age, sex, site and quality-controlled brain data included n=58,539 scans for log(JD) from a total of 52 scan sites, and n=53,593 scans for GMV/WMV from 50 scan sites. See Table 1 for demographic characteristics of the samples.

**Table 1:** Demographic characteristics of all samples. BL: baseline; 2Y: follow-up after two years; 4Y; follow-up after four years; SCA: Spinocerebellar ataxia (type 1 or 3).

| Datasets |  | N scans | N scan sites | Sex (F%/M%) | Mean age in years (SD) | Mean SARA score (SD) |
| --- | --- | --- | --- | --- | --- | --- |
| *Model estimation* |  |  |  |  |  |  |
| ABCD | Unique participants | 10,619 | 27 | 48.4/51.6 | - |  |
|  | age 9-11 | 9,846 | 26 | 48.7/51.3 | 10.0 (0.6) |  |
|  | age 11-13 | 6,946 | 26 | 46.9/53.1 | 12.0 (0.7) |  |
|  | age 13-15 | 1,259 | 23 | 47.3/52.7 | 14.1 (0.7) |  |
| CAMCAN |  | 593 | 1 | 50.9/49.1 | 54.6 (18.8) |  |
| CMI-HBN |  | 2,327 | 3 | 38.3/61.7 | 11.0 (3.4) |  |
| HCP-1200 |  | 1,111 | 1 | 54.5/45.5 | 28.8 (3.7) |  |
| HCP-Aging |  | 673 | 4 | 57.7/42.3 | 58.8 (14.9) |  |
| HCP-Dev |  | 643 | 4 | 51.2/48.8 | 13.8 (3.8) |  |
| IXI |  | 549 | 1 | 56.3/48.7 | 48.7 (16.4) |  |
| PING |  | 724 | 7 | 49.0/51.0 | 12.4 (5.0) |  |
| PNC |  | 1,369 | 1 | 50.8/49.2 | 14.2 (3.5) |  |
| UKB |  | 32,499 | 3 | 52.9/47.1 | 63.5 (7.6) |  |
| Total train set |  | 29,399 | 52 | 50.8/49.2 | 41.8 (26.1) |  |
| Total test set |  | 29,140 | 52 | 50.8/49.2 | 42.1 (26.1) |  |
| *Application to preterm birth* |  |  |  |  |  |  |
| genR | Unique subjects | 4736 | 2 | 50.9/49.1 | - |  |
|  | age 6-9 | 675 |  | 50.8/49.2 | 8.1 (1.0) |  |
|  | age 9-12 | 2997 |  | 50.9/49.1 | 10.1 (0.6) |  |
|  | age 13-15 | 2173 |  | 53.8/46.2 | 14.0 (0.6) |  |
|  | age 17-20 | 2327 |  | 51.7/48.3 | 18.6 (0.7) |  |
| genR preterm subset | Unique participants | 284 |  | 55.6/44.4 | - |  |
|  | age 6-9 | 40 |  | 47.5/52.5 | 7.9 (1.1) |  |
|  | age 9-12 | 168 |  | 54.8/45.2 | 10.2 (0.6) |  |
|  | age 13-15 | 137 |  | 58.4/41.6 | 14.1 (0.6) |  |
|  | age 17-20 | 148 |  | 50.0/50.0 | 18.7 (0.7) |  |
| ABCD preterm subset | Unique subjects | 304 | 26 | 48.7/51.3 | - |  |
|  | age 9-11 | 282 | 24 | 48.2/51.8 | 10.0 (0.6) |  |
|  | age 11-13 | 194 | 25 | 48.5/51.5 | 12.0 (0.6) |  |
|  | age 13-15 | 34 | 10 | 52.9/47.1 | 14.2 (0.6) |  |
| *Application to spinocerebellar ataxia* |  |  |  |  |  |  |
| Nijmegen | Controls | 20 | 1 | 40.0/60.0 | 50.2 (13.7) | - |
|  | SCA1 | 29 |  | 44.8/55.2 | 49.3 (13.6) | 12.3 (6.9) |
| Mexico | Controls | 12 | 1 | 41.7/58.3 | 44.0 (12.5) | - |
|  | SCA3 | 15 |  | 53.3/46.7 | 41.0 (12.7) | 13.5 (6.1) |

Finally, to ensure both computational feasibility and high spatial precision, we resampled log(JDs) brain maps to 2mm×2mm×2mm with continuous interpolation and masked them with the binarized mask of the whole-brain MNI152NLin2009cSym template (also resampled to 2mm×2mm×2mm), resulting in 235,818 voxels in total. We resampled GMV and WMV brain maps to 2mm×2mm×2mm, smoothed (FWHM = 6mm) and masked them with the thresholded binarized masks of the MNI152NLin2009cSym probabilistic tissue maps (probseg-GM and probseg-WM resampled to 2mm×2mm×2mm with probability > 0.5), resulting in 125,905 and 88,389 non-overlapping voxels in total, respectively.

### 2. Voxelwise normative models

#### 2.1. Model estimation

For each brain measure (log(JD), GMV or WMV), we estimated normative models for each voxel with the warped Bayesian Linear Regression (BLR) approach from the PCNtoolkit (https://github.com/amarquand/PCNtoolkit/, version 1.1.2) in Python (version 3.12). We used age as a covariate, with a B-spline basis expansion to model non-linear effects as in prior work^45^ and binary sex and site as batch effects, with a train-test split of 50-50 stratified by site and sex. As the ABCD cohort includes multiple scans per participant, we also stratified this dataset such that same-participant scans would end together into either train or test set, not split between both.

The models thus produced site-corrected, sex- and age-specific normative deviation scores (z-scores) reflecting the degree to which brain anatomy deviated from the biological norm at voxel-level^8,9,46^. Fit metrics included explained variance, as well as excess kurtosis and skewness which measure tail-heaviness and asymmetry of the z-scores distribution, respectively, as used previously^11^.

Using the trained normative models, we analyzed two sets of participants born preterm, both with longitudinal MRI data. The first set was a subsample from ABCD (held-out from model training) and included participants with preterm birth (n=304 unique participants) based on the question ‘how many weeks premature was the child when they were born?’ from the abcd_dev_hxss01.txt questionnaire. Following Thalhammer et al. ^30^, we included participants using a strict threshold of gestational age ≤ 32 weeks, to reduce misclassification due to the potentially noisy, retrospective recall of gestational age in that measure. All ABCD participants with gestational age ≤ 37 weeks (i.e. both the participants included in preterm analyses, and the participants with less certain prematurity) were explicitly retained in the test set during model estimation, such that they remained fully held-out from the model training set.

We then transferred the estimated models to an independent sample, Generation R (GenR)^47,48^, a prospective, population-based cohort from fetal life onwards. Of note, the aggregated reference normative data and the GenR data were hosted in different institutions, but the transfer to GenR required only the sharing of trained models parameters, precluding the need for any outgoing GenR data transfer.

Longitudinal MRI scans of GenR participants were collected across four timepoints every 3-4 years (age 6-9, age 9-12, age 13-15, age 17-20) and 2 scanners. (see supplementary material for MRI acquisition parameters). We conducted preprocessing of GenR T1 scans as was described in the previous section (ANTS version 2.5.4), and final quality-controlled sample size included n=8,172 scans (n=4736 unique participants, see Table 1). We used a calibration subsample (five percent of scans, n=493, stratified by batch effects with the rest of the GenR sample) with the BLR transfer function of the PCNtoolkit to estimate new site effects and adjust model predictions accordingly.

The GenR dataset thus provided the second set of participants born preterm (n=284 unique participants with gestational age ≤ 37 weeks measured at birth). We explicitly kept these participants out of the calibration set during model transfer. Please see Table 1 for the demographic characteristics of both ABCD and GenR preterm subsets.

In both ABCD and GenR, we compared voxelwise differences in the proportion of extreme z-scores (|z-scores| ≥ 2) at each timepoint between participants with preterm birth and a same-sized group of controls born at term randomly selected from the same dataset. For each comparison, we further assessed the spatial stability of results over five non-overlapping control subgroups, randomly selected from the same dataset. We assessed group differences in the total brain burden of extreme z-scores (i.e. the percentage of voxels with extreme z-score relative to the total number of voxels) at each timepoint with Mann-Whitney U-tests and Bonferroni-Holm correction^49^ to correct for multiple testing across timepoints and directions (positive and negative extreme deviations), using all available controls. We evaluated mean effects group differences at each timepoint by comparing normative z-score brain maps with threshold-free cluster enhancement (TFCE) in nilearn with 5000 permutations, and we report clusters with FWE-corrected p-values<0.05. For the TFCE analysis each group of controls was set to ten times the number of participants with preterm birth, as a compromise between leveraging the available statistical power and keeping reasonable computational costs of permutation testing. Because GMV and WMV normative deviation maps had no overlapping voxels, they were merged into a single whole-brain deviation map per participant, where grey matter voxels showing deviations from the GMV normative models and white matter voxels showing deviations from the WMV models.

#### 2.3. Application to spinocerebellar ataxia type 1 and 3

We also transferred the estimated voxelwise normative models to small clinical samples of patients with spinocerebellar ataxia (SCA) and healthy age-matched controls from two independent studies, with data collected in Nijmegen^50^ and Mexico^51^. Participant recruitment and MRI acquisition parameters are described in the supplementary material. Briefly, all SCA patients were assessed with the Scale of Assessment and Rating of Ataxia (SARA), which is a 40-point scale assessing ataxia symptoms^52^ including difficulties with gait, sitting, posture, speech, and finger and hand movements. Patients from Nijmegen were re-assessed a year later. Final sample sizes for log(JDs) analyses included n=29 SCA1 patients and n=20 controls from Nijmegen, and n=15 SCA3 patients and n=12 controls from Mexico (see Table 1). Two more patients from Nijmegen and three patients from Mexico were excluded for GMV/WMV analyses due to segmentation failures. Each SCA sample was processed separately, and preprocessing of MRI data was conducted as described above (ANTs 2.2).

For each SCA sample, we used healthy controls from the same scan site as calibration set to adjust normative models with new scan site effects. We then calculated whole-brain voxelwise z-scores for each patient, reflecting individual morphometric deviations from the biological norm, and we extracted brain maps of extreme z-scores (|z-scores| ≥ 2). Here too, GMV and WMV normative deviations were combined into a single normative deviation map per participant.

Additionally, we conducted exploratory symptom severity prediction analyses using scikit-learn (version 1.7.1). We used four sets of brain measures at baseline (z-scores or raw measures, log(JD) or combined GMV and WMV) as feature sets in separate regression analyses to predict ataxia severity (SARA score) at baseline, and severity score change from baseline to follow-up. We used a kernel ridge regression (KRR) model^53^, a multivariate regularized regression approach. The use of a linear kernel space allows to keep a low computational cost as computational complexity scales with sample size and not with feature dimensionality, which is desirable in analyses that include whole-brain voxelwise features.

We kept the complexity of the predictive pipeline minimal, in consideration of the very small sample sizes. We used default parameters of the KRR regressor, adding an explicit bias term (column of ones) to account for the mean. We also compared the deviations from the normative models to raw features, which were first age-residualized and standardized. Given the sample sizes, we used a leave-one-out cross-validation approach. We first fit the KRR using only the Nijmegen sample as a training set, then using pooled Nijmegen and Mexico samples (the Mexico sample was too small to be a standalone testing set), with all other pipeline parameters kept identical.

## Results

### 1. Voxelwise normative models provide good fit to the data

First, we demonstrated that voxelwise normative models provided an excellent fit to the data. Specifically, models fit with age as covariate and sex and site as batch effects, accounted for as much as 78%, 85% and 82% of the variance in log(JD), GMV and WMV respectively. Overall, this pattern recapitulated known trajectories of brain development at high resolution^7,54^ (see Fig. 2 for explained variances for log(JDs) models, Fig. 3 for explained variance in GMV and WMV models). For example, explained variance was generally higher in the ventricles and in the surrounding white matter. Importantly, across all modalities, the shape of the modeled distribution was appropriate for the data in that the z-score distributions derived from the voxelwise models generally showed acceptably low skewness (|skew| < 1) and low excess kurtosis (|kurtosis|<5) (see supplementary Fig. 1 for kurtosis, skewness and additional fit metrics in both sets of models).

**Figure 2:**
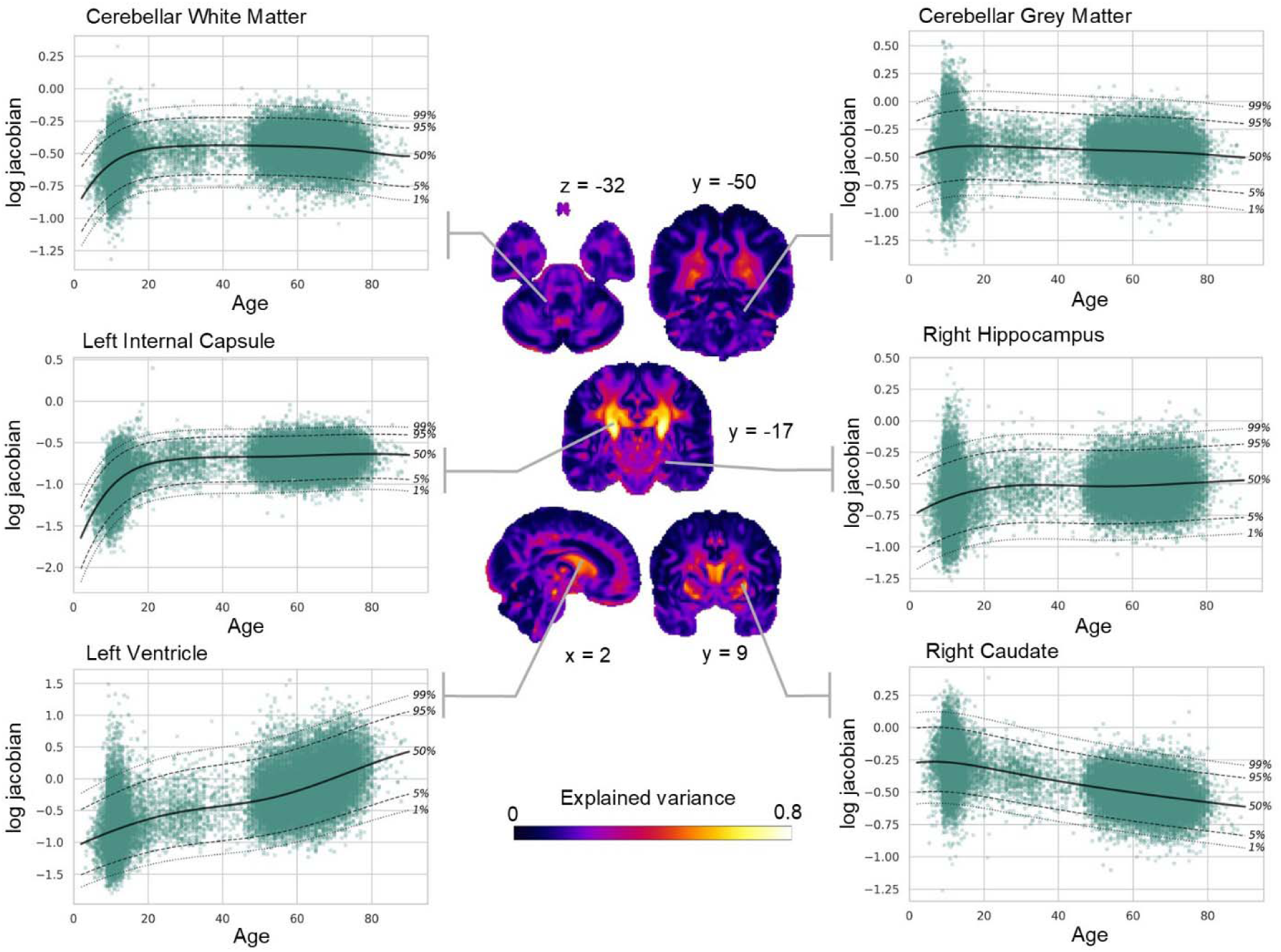
Voxelwise normative model fit of log jacobian determinants across the lifespan. Explained variance of the log jacobian determinant model on the test set is shown, alongside normative trajectories harmonized for batch effect (sex and site) from example voxels.

**Figure 3:**
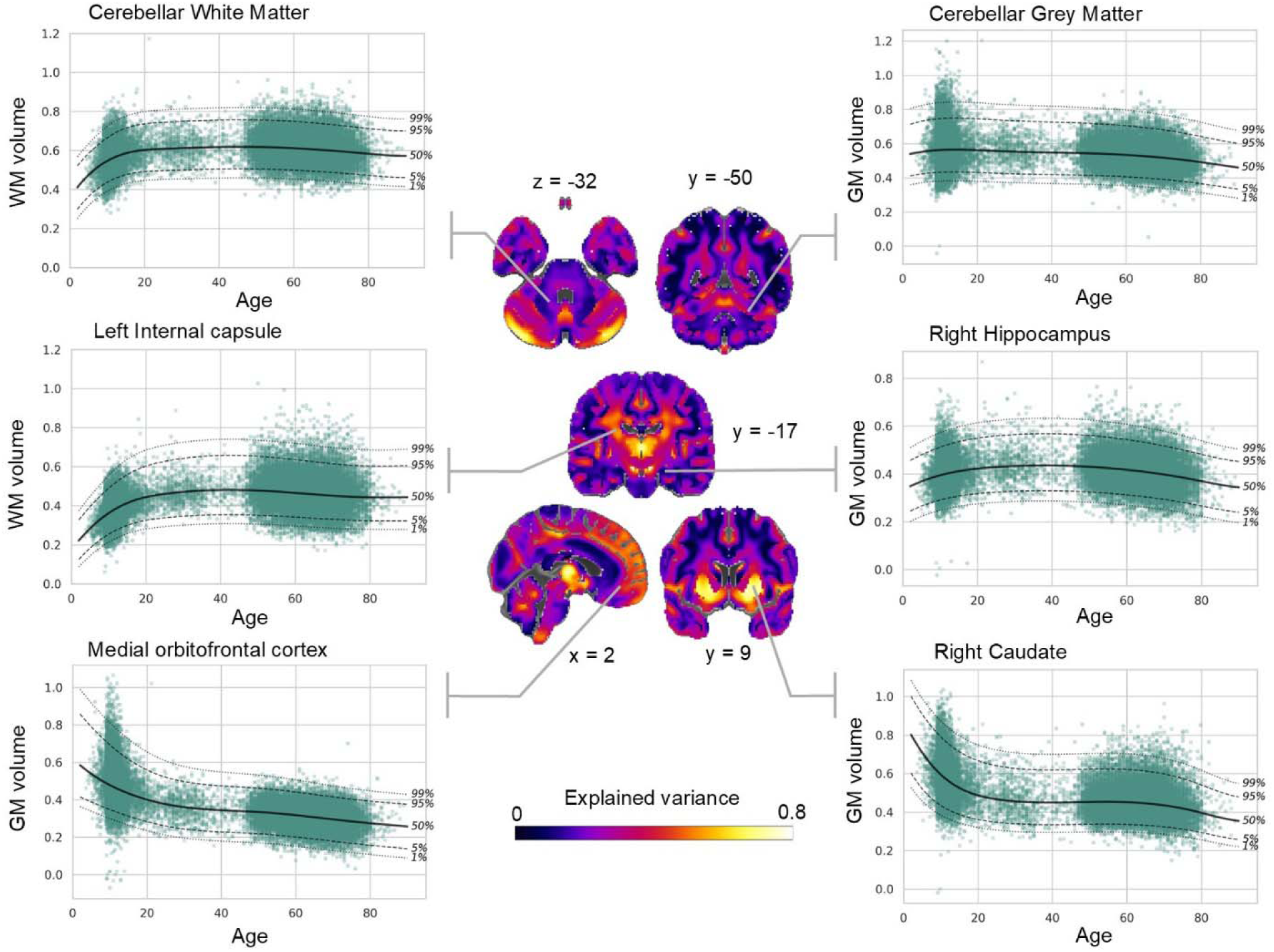
Voxelwise normative model fit of grey and white matter volume across the lifespan. Explained variance of the grey matter and white matter models on the test set is shown, alongside normative trajectories harmonized for batch effect (sex and site) from example voxels. As grey and white matter models were estimated on non-overlapping voxels, both modalities are displayed in a combined, whole-brain approach. GM: grey matter; WM: white matter.

### 2. Preterm birth gives rise to widespread and persistent patterns of deviation

Next, we applied this model to preterm-born individuals in two comparable youth cohorts (ABCD and Generation R). In the ABCD subsample, which was held out from model training, a higher proportion of participants born preterm showed extreme negative z-scores relative to controls born at-term in a widespread pattern of voxels across the brain using log(JDs) normative models (see Fig. 4A for log(JDs) extranormal z-scores and supplementary Fig. 2A for GMV-WMV extranormal z-scores).

**Figure 4:**
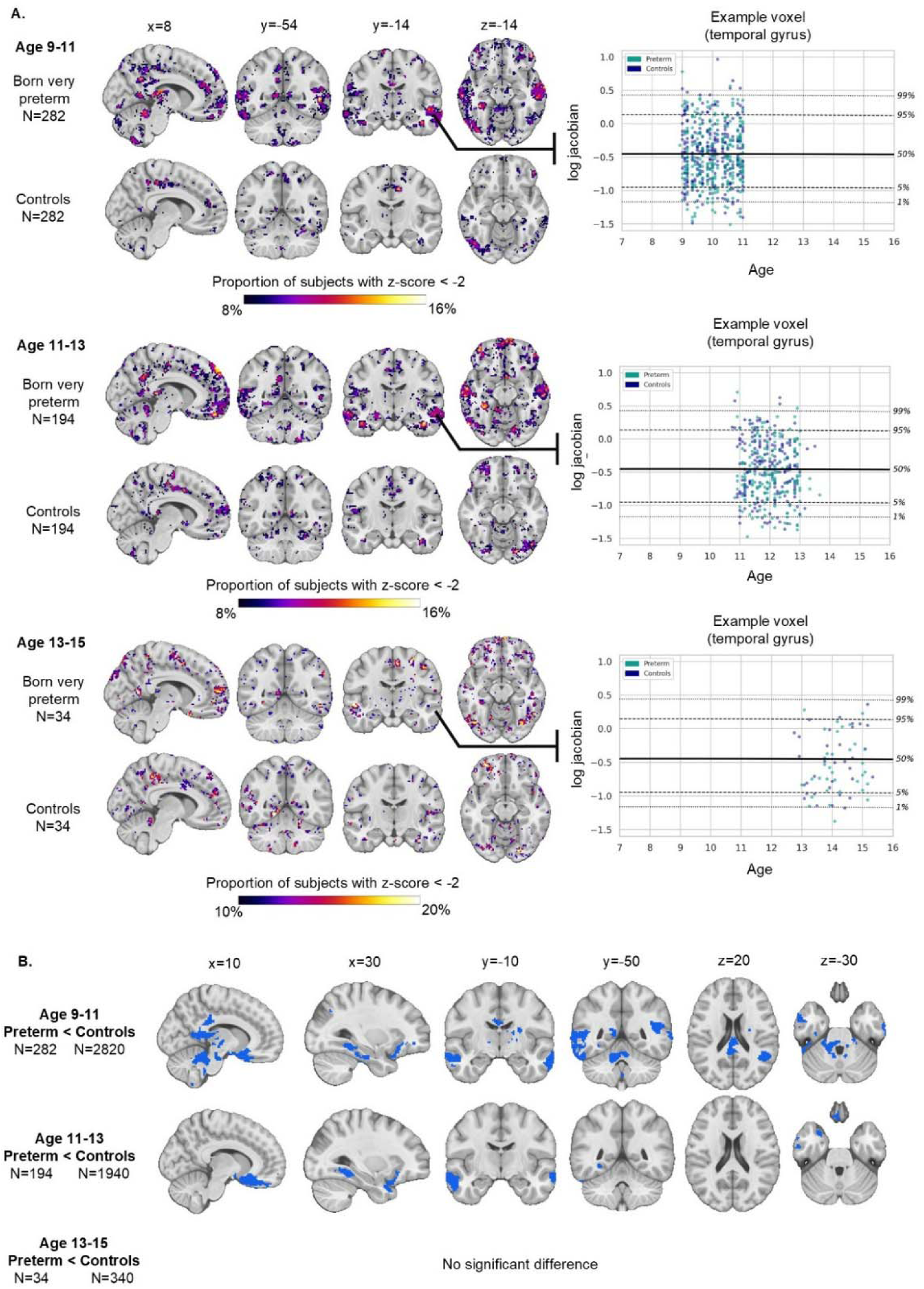
Normative deviations of log jacobian determinants after preterm birth in the ABCD dataset. A: Voxelwise proportions of extranormal negative deviations (z-score<-2) in ABCD participants born preterm compared to a same-sized group of controls born at-term at ages 9-10, 11-13 and 13-15. Normative trajectories harmonized for batch effect (sex and site) are shown for example voxels; B: Voxelwise mean z-score differences between participants born preterm and a tenfold control group born at-term, at all three timepoints. Suprathreshold clusters with TFCE p<0.05 FWE-corrected are shown.

This voxelwise difference in proportion of extreme negative deviations persisted across the first two timepoints (ages 9-11 and 11-13), and remained stable over five randomly sampled same-sized control subgroups. The total brain burden of extremely negative deviations (i.e. the number of voxels with extreme deviation over the total number of voxels) was also subtly but significantly larger in participants born preterm compared to controls (Holm-corrected p=0.041 at both timepoints). Voxelwise differences in the proportion of extreme negative z-scores between the two groups using GMV-WMV models at ages 9-11 and 11-13 were less distinct, but still visible in the cerebellum, thalamus and precuneus. However, we did not detect significant differences in total brain burden of extreme deviations for GMV/WMV models between groups at any timepoint. We also found no voxelwise proportion differences nor total brain burden differences in extreme positive deviations between participants born preterm and controls for any set of models. It is important to note that the overlap across participants for any given voxel remained relatively low (<20% of participants), which is a reflection of the degree of inter-individual variability in this cohort.

Next, we aimed to determine which regions showed group-level mean effects of preterm birth. We detected clusters of more negative z-scores in participants born preterm compared to controls at age 9-11 in widespread areas across the brain, including but not limited to the brainstem, cerebellum, bilateral temporal cortices, splenium of corpus callosum and medial prefrontal cortex using log(JDs) models (p < 0.05 FWE; Fig. 4B and supplementary Fig. 2B). At age 11-13, we found significant clusters of more negative deviations in the medial prefrontal cortex and bilateral temporal cortices in participants born preterm. We only detected significant group differences in z-scores from the GMV-WMV models at age 11-13, in clusters in the left temporal, bilateral medial prefrontal, and right inferior frontal cortices. We did not find any significant cluster with more positive deviations in participants born preterm compared to controls at any timepoint, for both sets of models.

Next, we aimed to replicate this finding in an independent dataset (GenR) which, like the ABCD subsample above, was not part of the normative model training procedure. In this dataset, a higher proportion of participants born preterm showed extreme negative z-scores relative to controls born at-term in a more widespread pattern of voxels across the brain, including but not limited to the brainstem, cerebellum, bilateral temporal cortices, splenium of corpus callosum, internal capsule and medial prefrontal cortex (see Fig. 5A for log(JDs) extranormal z-scores and supplementary Fig. 3 for GMV-WMV extranormal z-scores). This voxelwise difference in the proportion of extreme negative deviations was less distinct at age 6-9, but was very consistent across the three other timepoints (ages 9-12, 13-15, 17-20) and across both log(JDs) and GMV-WMV models, although less distinct for the latter. Results remained stable over five randomly sampled same-sized control subgroups and again the level of overlap across participants was relatively low for any given voxel (<15%). Across these three timepoints, the total brain burden of extremely negative deviations was larger in participants born preterm compared to controls (Holm-corrected p=4.6e-4 at age 9-12, p=0.016 at age 13-15, p=1.8e-4 at age 17-20 for log(JDs) deviations; p=1.7e-4 at age 9-12, p=0.031 at age 13-15, p=1.9e-4 at age 17-20 for GMV-WMV deviations), but no significant difference was found at age 6-9. We found no visible voxelwise differences nor total brain burden differences in extreme positive deviations between participants born preterm and controls.

**Figure 5:**
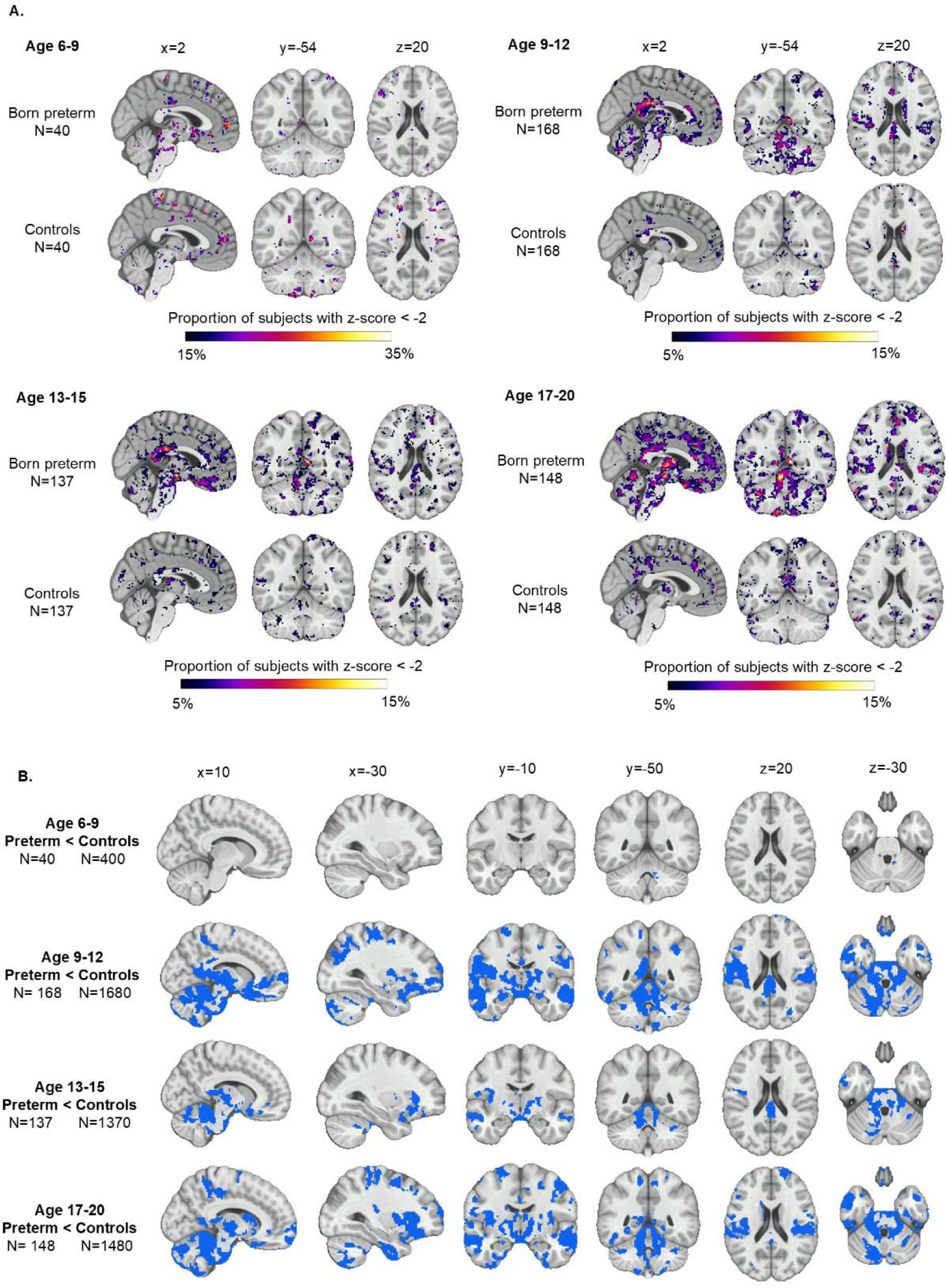
Normative deviations of log jacobian determinants after preterm birth in the GenR dataset. A: Voxelwise proportions of extranormal negative deviations (z-score<-2) in GenR participants born preterm compared to a same-sized group of controls born at-term at ages 6-9, 9-12, 13-15 and 17-20. Normative trajectories harmonized for batch effect (sex and site) are shown for example voxels; B: Voxelwise mean z-score differences between participants born preterm and a tenfold control group born at-term, at all four timepoints. Suprathreshold clusters with TFCE p<0.05 FWE-corrected are shown.

A group level analysis of mean effect differences again revealed significant clusters of more negative z-scores in participants born preterm compared to controls at ages 9-12, 13-15, 17-20 in a widespread pattern of voxels across the brain, including but not limited to the brainstem, cerebellum, bilateral temporal cortices, splenium of corpus callosum, internal capsule and medial prefrontal cortex (see Fig. 5B and supplementary Fig. 3B). This was the case for both log(JDs) and GMV-WMV models. At age 6-9, we detected fewer but still significant clusters in the brainstem, cerebellum and putamen in GMV-WMV models, and only a small cluster in the cerebellar white matter was significant in the log(JDs) models. We did not find any significant cluster of more positive z-scores in participants born preterm compared to controls at any timepoint.

### 3. SCA individual-level deviation maps show distinct heterogeneity

Next, we aimed to demonstrate the clinical utility of our voxelwise models for parsing heterogeneity in spinocerebellar ataxia. Patients with spinocerebellar ataxia presented with marked neurobiological heterogeneity; in Figure 6 we present examples of patient-level maps of voxelwise extranormal normative deviations from patients with spinocerebellar ataxia type 1 and 3 (maps for the full patient samples are included in supplementary Fig. 4 and 5 for the Nijmegen SCA1 and Mexico SCA3 studies respectively). These brain maps illustrated considerable variability in the neurobiological effects of each genetic mutation. Whilst most patients showed evidence of atrophy in the pons or brainstem, effects were divergent in other regions: cerebellar atrophy was only evident for some individuals, and some individuals showed increased volume in the ventricles, whilst others showed widespread cortical effects. This once again underscored the need for parsing heterogeneity at high spatial resolution. Also, it is important to note that the degree of atrophy was only partially related to symptom severity. For instance, some individuals had widespread atrophy, yet were not severely impaired according to the SARA measure.

**Figure 6:**
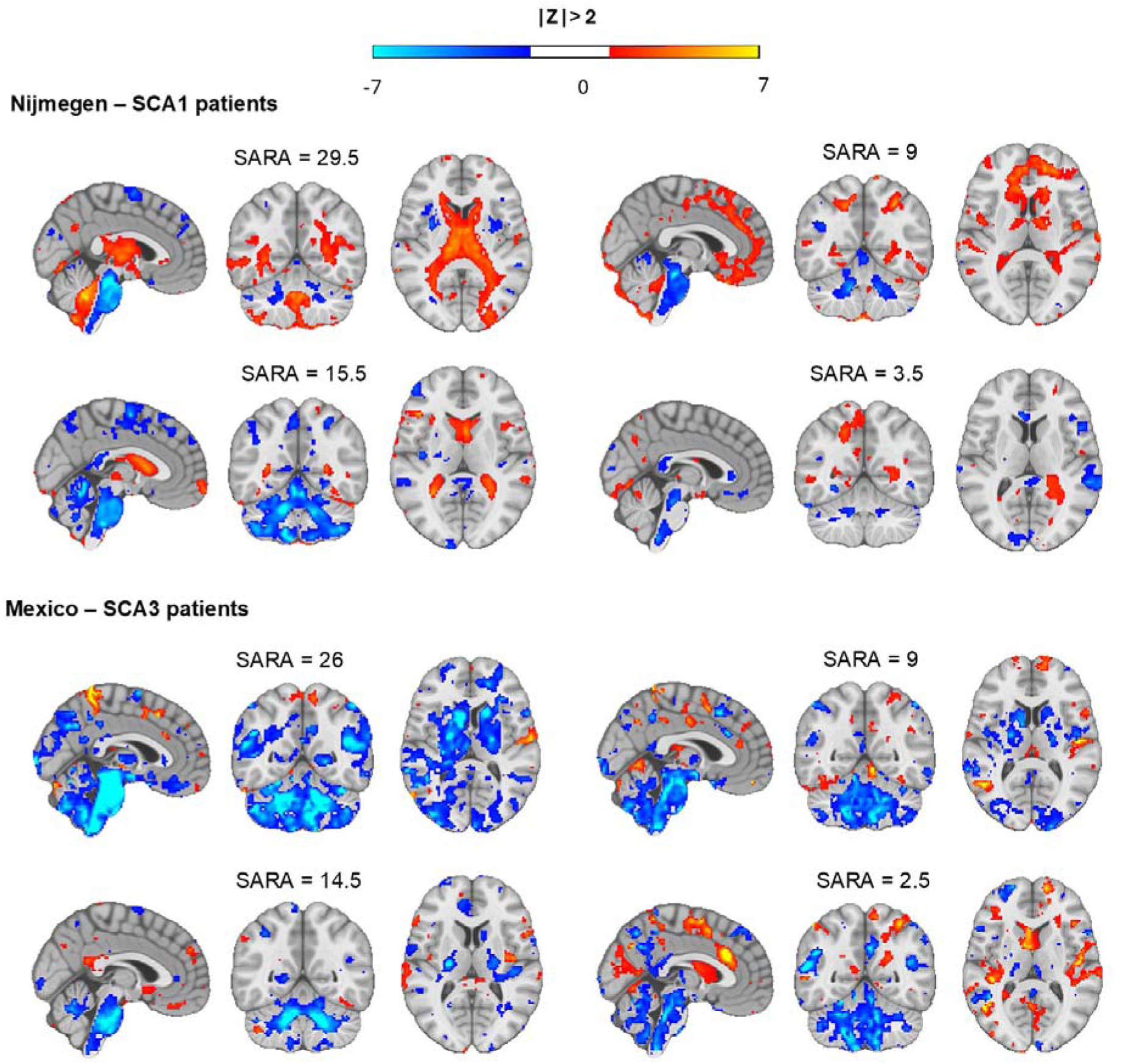
Example whole-brain maps of log jacobian determinants extranormal deviations (z-scores) in patients with spinocerebellar ataxia type 1 and 3. SCA: spinocerebellar ataxia (type 1 or 3); SARA: Scale for the Assessment and Rating of Ataxia.

Next, we used these deviation maps for exploratory prediction of ataxia symptom severity using a multivariate kernel ridge regression approach. This yielded moderate to good predictive performance within the Nijmegen SCA1 sample using leave-one-out cross-validation (explained variance/r² ranging from 0.18 to 0.65, see Table 2 for all prediction results), with combined GMV and WMV measures yielding better performance than log(JDs) measures, and normative measures yielding better prediction performance than the corresponding raw measures. Using a pooled sample of Nijmegen SCA1 and Mexico SCA3 samples with leave-one-out cross validation yielded similar predictive performance (explained variance/r² ranging from 0.18 to 0.58). Voxelwise training feature weights were widespread across the brain but consistently higher in the brainstem and cerebellar grey and white matter across all measures (see supplementary Fig.6). The prediction of prospective clinical severity change after one-year follow-up using on baseline brain measures yielded low predictive accuracy within the Nijmegen SCA1 sample with leave-one-out cross-validation (r² ranging from 0.13 to 0.18).

**Table 2.** : Clinical severity prediction results in spinocerebellar ataxia type 1 and 3. LOOCV: Leave-one-out cross validation; SCA: Spinocerebellar ataxia type 1 or 3; SARA: Scale for the assessment and rating of ataxia.

| Feature sets | LOOCV<br>Nijmegen<br>(SCA1)<br>$r^2$ | LOOCV<br>pooled Nijmegen (SCA1) +<br>Mexico (SCA3)<br>$r^2$ |
| --- | --- | --- |
| *Prediction of baseline clinical severity (SARA score)* |  |  |
| Log jacobian determinants raw | 0.18 | 0.18 |
| Log jacobian determinants Z-scores | 0.40 | 0.35 |
| Grey and white matter volumes raw | 0.57 | 0.43 |
| Grey and white matter volumes Z-scores | 0.65 | 0.58 |
| *Prediction of clinical severity change between baseline and follow-up (SARA slope)* |  |  |
| Log jacobian determinants raw | 0.17 |  |
| Log jacobian determinants Z-scores | 0.13 |  |
| Grey and white matter volumes raw | 0.18 |  |
| Grey and white matter volumes Z-scores | 0.15 |  |

## Discussion

In this paper, we estimated normative models of voxel-level volumetric measures for the whole brain, across the lifespan based on a large, aggregated reference sample and using a state-of-the art image preprocessing pipeline. We showcased their utility in two clinical applications: long-term effects of preterm birth, and patient-level alterations in a rare neurodegenerative disease (spinocerebellar ataxia). In both cases, we show clinically relevant effects and inter-individual variability evident at a considerably finer granularity than can be readily studied using classical region-level analysis. To maximize the utility of this contribution we distribute these models freely to the community accompanied by a set of open-source software tools. We additionally make available GMV and WMV normative models with larger brain coverage (estimated using tissue map probability > 0.2 with all other parameters kept identical).

In our first clinical application, we show that a larger proportion of participants born preterm had extreme negative volumetric deviations (z-score < −2) from the sex-, site- and age-specific biological norm, in voxels widespread across the brain, compared to controls. Negative deviations correspond to either volume contraction (log(JDs)) or reduced tissue volume (combined GMV and WMV) relative to the mean and variance in the reference normative model. These effects were distributed across widespread areas and in both ABCD and GenR cohorts in a generally temporally consistent manner, spanning ages 9 to 20, but not at age 13-15 in ABCD nor age 6-9 in GenR. However, this can be explained by the substantially reduced sample size at each of these timepoints (approximately 70% and 80% reduction in the number of preterm individuals in GenR and ABCD respectively). We also showed spatially distributed mean group effects, as clusters of significantly more negative deviations were found in participants born preterm compared to controls born at-term in both cohorts and for most timepoints. However, heterogeneity across participants was still very distinct in the normative deviation brain maps, and the proportion of participants born preterm with extreme deviations very rarely exceeded 20% for any voxel, model and timepoint. This underscores the need for mapping normative deviations at a high spatial resolution.

In our second clinical application, we produced brain maps of extreme normative deviations for each SCA1 and SCA3 patient. These normative deviation brain maps were in line with well-documented pontine and cerebellar atrophy for both SCA1 and SCA3^31–34^, but they also illustrated striking inter-individual heterogeneity in the neurobiological disease effects that only showed a partial correspondence with symptom levels. This further underscores the value of generating statistical disease fingerprints from normative models that can be used for personalized inference.

We conducted an exploratory multivariate prediction of cross-sectional ataxia severity that yielded accurate cross-validated prediction performance within the Nijmegen SCA1 sample and within the pooled sample of Nijmegen and Mexico (SCA1 and SCA3 respectively). Normative measures performed better than raw features for this cross-sectional analysis. We also conducted a prediction analysis of prospective disease progression at one-year follow up in the Nijmegen sample using the same measures, but predictive performances were low for all measures. This indicates that cross-sectional volumetric measures at baseline are not predictive of outcome. Note that whilst we previously reported significant associations of regional pontine and cerebellar volumes with disease progression in the same Nijmegen SCA1 sample, this apparent discrepancy can be explained by the more stringent, out-of-sample prediction strategy that was used in the present work^34^.

Both clinical applications highlighted distinct but complementary benefits of our voxelwise normative modeling approach. The first and more general aspect is that normative models are able to detect individual-level signals of interest in voxel-level brain measures that are not necessarily captured by group-level mean differences. Normative deviations can then be used in downstream analyses as biological variables that cleanly account for covariates such as sex and scanning site. Here, while we show comparisons of normative deviations between groups using participants born preterm and participants born at-term, we also illustrate the individual-level heterogeneity underlying this group contrast. Associations between normative deviations and relevant behavioral traits or clinical outcomes can also be investigated at the individual level using less categorical approaches such as, for example, regression or soft clustering techniques^9,10^. As such, the present work shows that normative modeling scaled to voxel precision can be a useful tool to move beyond group-level case-control analyses and towards individualized profiles of disease, resilience, risk of onset or relapse.

The second benefit illustrated by the present work is that normative models can provide individual-level, voxelwise maps of extreme normative deviations, reflecting potential brain alterations for each patient. By leveraging a large reference cohort, normative models can be transferred to clinical samples from new sites, provided sufficient calibration samples are used. This can be of particular interest for rare disorders with small clinical sample sizes, as we show with spinocerebellar ataxia. Cross-referencing these voxel-level deviation maps with, for instance, neurotransmitter density or gene expression brain maps, could be a promising step towards sample stratification or enrichment in clinical trials for treatments targeting specific pathways in the long term.

Several limitations of our normative models should be noted. First, individuals from western, educated, industrialized, rich and democratic countries are overrepresented in available large-scale datasets and thus our models, trained with such a reference sample, inaccurately represent the general population and might not generalize to different populations^55,56^. Although our aggregated sample has reasonable coverage of the lifespan, with the exception of neonates and infants, one should be careful in interpreting normative deviations where training data is more sparse, for example the edges of the age distribution, as it leads to larger uncertainty in the estimated centiles^57^. Extrapolating beyond the training age range should be avoided, although models may be transferred using a model re-estimation step to different age ranges if necessary^58^. Finally, our normative models should be transferred to new data only if these data were extracted with the same MRI preprocessing pipeline that was used in training. To this end, we have kept most default parameters of the ANTS and fsl_anat preprocessing pipelines, so that they are maximally transferrable and we freely share our code for preprocessing and normative models transfer.

Second, in the application to preterm birth, we had limited data for individuals aged 13-15 in ABCD and 7-9 in GenR, and thus associated results should be interpreted as exploratory at this time, although additional validations for the ABCD sample can be performed as additional data are released.

Third, in the ABCD dataset, the degree of prematurity was only measured retrospectively by asking parents 10 years after childbirth, leading to noisy measurement of gestational age and, importantly, ABCD explicitly excludes individuals with extremely preterm births (<28 weeks) as well as several associated medical conditions such as brain hemorrhage or cerebral palsy, potentially resulting in a less affected preterm sample.

Finally, in the application to spinocerebellar ataxia, given the very low prevalence of SCA1 and SCA3, and while the Nijmegen SCA1 sample is one of the largest single-center SCA1 samples to date, clinical sample sizes were still small and only allowed for exploratory predictions. To accommodate this, we employed a penalized regression approach and kept predictive pipeline complexity to a minimum.

For individuals born preterm, a crucial next step in investigating neuroanatomical heterogeneity would be to extend the age coverage of voxel-level normative models to infants and neonates. This would enable the investigation of the associations between individual-level neuroanatomical normative deviations in the first weeks or months of life and prospective neurodevelopmental outcomes. Among possible research questions, this could help identify risk factors and improve the accuracy of existing predictive models, with potential to ultimately facilitate targeted monitoring and medical care^59^. It is important to note, however, that serious challenges in neuroimaging data acquisition and processing remain in that age range^60^, and that further methodological fine-tuning might also be necessary to be compatible with a scalable, normative modeling approach^61^.

For rare diseases such as spinocerebellar ataxia, collective efforts such as ENIGMA-ataxia ((http://enigma.ini.usc.edu/ongoing/enigma-ataxia/) are especially important. Beyond a limited increase in clinical sample size, however, collecting sufficient control data is also essential to be able to accurately adjust normative models for new sites and fully leverage the normative modeling framework even for small patient samples. However, it is important to reiterate that the normative modelling approach allows us to move beyond group-level inferences, and can be used to study atypicalities in individual participants. For example, to understand the effects of highly penetrant, rare genetic polymorphisms on the brain^20^.

Finally, an exciting perspective for future research using neuroanatomical normative modeling is to study change over time using longitudinal data^62,63^. Indeed, the way individual-level normative deviations change over time could reflect clinically meaningful information, and it is especially useful to be able to do this at high spatial resolution. This can enable the detection of individual deviations in rates of change in longitudinal neuroimaging datasets, which could for instance be promising to investigate in pre-symptomatic mutation carriers beginning to deviate from the reference population^64^. Overall, in this work, we demonstrate a scalable normative modelling approach at voxel-level precision using structural MRI, allowing us to parse voxelwise neuroanatomical heterogeneity at the individual level, and we illustrate this approach with two use cases, namely mapping the neurobiological sequelae of preterm birth and understanding inter-individual variability in neurodegenerative disease. In order to maximize the utility of this contribution, we freely distribute all models to the community.

## Conflicts of interest

The authors report no biomedical commercial relationships or conflicts of interest.

## Acknowledgements

This work had been supported by a funding grant from the Raynor Cerebellum Project and has been conducted using the UK Biobank resource under application number 23668. RLM was also supported by the European Union’s HorizonEurope Research and Innovation Programme (FAMILY, grant agreement No 101057529). The Nijmegen SCA1 study was supported by a grant from ZonMw (40-44600-98-606).

## Notes

### Competing Interest Statement

The authors have declared no competing interest.

