## Supplementary methods and figure legends for "Dissecting heterogeneous brain development and aging using voxelwise normative models"

1. GenR MRI scan acquisition

For the Generation R sample^1,2^, scans were acquired using two different MRI scanners. In the first timepoint visit, data were collected on a GE MR750 Discovery system. High resolution T_1_- weighted MRI scans were acquired using an inversion recovery fast spoiled gradient recalled sequence (IR-FSPGR) with T_R_ = 10.3 ms, T_E_ = 4.2 ms, T_I_ = 350 ms, flip angle = 16°, acquisition time = 5 min 40 s, field of view = 230.4 × 230.4 mm, 0.9 × 0.9 × 0.9 mm^3^ isotropic resolution. Images were resampled to 1 mm isotropic resolution to match data from the subsequent timepoints.

In the second, third and fourth timepoints, data were collected on a study-dedicated GE MR750w system (General Electric Healthcare, Wisconsin, USA). High resolution T_1_- weighted MRI scans were acquired using an IR-FSPGR sequence with T_R_ = 8.77 ms, T_E_ = 3.4 ms, T_I_ = 600 ms, flip angle = 10°, acquisition time = 5 min 20 s, field of view = 220 ×220 mm, 1x1x1 mm^3^ isotropic resolution.

2. SCA samples and MRI scan acquistion

For the Nijmegen SCA1 sample^3^, patients were recruited between 2020 and 2022 via patient associations and the outpatient clinic of the Radboud UMC Medical Center. Healthy controls were recruited in parallel and consisted mostly of partners or non-affected family members of patients. All participants were ≥ 18 years-old and had no other neurological disease. Criteria for patients with SCA1 to be included were a positive molecular genetic test (pathogenic CAG repeats expansion in the ATNXN1 gene) and a SARA total score of ≥ 3 at any of the three assessment visits, thus allowing pre-symptomatic inclusion at baseline. MRI scanning and clinical assessment, including SARA scale, were conducted at baseline, one-year follow-up, and two-year follow-up.

Neuroimaging data for the Nijmegen SCA1 sample were acquired using a 3T MRI scanner (Magnetom Prisma-fit, Siemens Healthineers, Erlangen, Germany). The acquisition protocol included a high-resolution 3D inversion-prepared T1-weighted image (TR = 2.3ms, TE = 4.48ms, TI = 950ms, 0.9 mm isotropic) and an MRS package developed by Drs Öz and Deelchand provided by the University of Minnesota.

For the Mexico SCA3 sample^4^, patients were recruited on the basis of a molecular diagnosis (abnormal CAG repeats expansion in the ATNXN3 gene). The control group consisted in age- and gender-matched healthy volunteers. Inclusion criteria for patients and controls were no history of neurological disease other than SCA and no neuropharmacological treatment.

For the Mexico SCA3 sample, MRI data were acquired using a 32-channel quadrature head coil in a 3T Achieva MRI scanner (Phillips Medical Systems, Eindhoven, The Netherlands). The acquisition protocol included a high-resolution 3D T1 Fast Field-Echo sequence, with TR/TE of 8/3.7 ms, FOV of 256 × 256 mm, flip angle 25° and an acquisition and reconstruction matrix of 256 × 256, resulting in an isometric resolution of 1 × 1 × 1 mm^3^.

All patient recruitments were approved by the respective local ethics committees (Nijmegen: CMO-2019-5377; Mexico: ethics committees on human experimentation of the Universidad Nacional Autonoma de Mexico) and written informed consent was obtained for all participants prior to participation in the respective studies.

Supplementary figure legends

**Supplementary Figure 1: Additional fit metrics of voxelwise normative models.** A : Fit metrics for log jacobian determinants models; B: Fit metrics of grey and white matter volume models. MACE : Mean absolute scaled error; MSLL : Mean standardized log loss. As grey and white matter models were estimated on non-overlapping voxels, both modalities are displayed in a combined, whole-brain approach.

**Supplementary Figure 2:** **Normative deviations of grey and white matter volumes after preterm birth in the ABCD dataset.** A: Voxelwise proportions extranormal negative deviations (z-score<-2) in ABCD participants born preterm compared to a same-sized group of controls born at-term , at ages 9-11, 11-13 and 13-15. Normative trajectories harmonized for batch effect (sex and site) are shown for example voxels; B: Voxelwise mean z-score differences between participants born preterm and all controls, at all three timepoints. Suprathreshold clusters with TFCE p<0.05 FWE-corrected are shown. As grey and white matter models were estimated on non-overlapping voxels, both modalities are displayed in a combined, whole-brain approach.

**Supplementary Figure 3:** **Normative deviations of grey and white matter volumes after preterm birth in the GenR dataset.** A: Voxelwise proportions of extranormal negative deviations (z-score<-2) in GenR participants born preterm compared to a same-sized group of controls born at-term at ages 6-9, 9-12, 13-15 and 17-20. Normative trajectories harmonized for batch effect (sex and site) are shown for example voxels ; B: Voxelwise mean z-score differences between participants born preterm and a tenfold control group born at-term, at all four timepoints. Suprathreshold clusters with TFCE p<0.05 FWE-corrected are shown. As grey and white matter models were estimated on non-overlapping voxels, both modalities are displayed in a combined, whole-brain approach.

**Supplementary Figure 4:** **Whole-brain maps of log jacobian determinants extranormal deviations (|z-score|>2) in patients with spinocerebellar ataxia type 1 from the Nijmegen study.** SARA: Scale for the Assessment and Rating of Ataxia; BL: baseline; Y1 : one-year follow-up.

**Supplementary Figure 5:** **Whole-brain maps of log jacobian determinants extranormal deviations (|z-score|>2) in patients with spinocerebellar ataxia type 3 from the Mexico study.** SARA: Scale for the Assessment and Rating of Ataxia.

**Supplementary Figure 6: Voxelwise feature training weights from the predictions of baseline clinical severity in Spinocerebellar ataxia type 1 within the Nijmegen sample.** Feature weights were averaged across all leave-one-out training sets.
